# Mechanistic Analyses of Supercoiling Behaviors of DNA in Nucleosomes and Chromatins

**DOI:** 10.1101/523084

**Authors:** Hao Zhang, Tianhu Li

## Abstract

Besides those in 146-base pair nucleosome core particle DNA, supercoils have been known to be present in 10-base pair arm DNA segments and naked linker DNA segments. The interacting patterns among histone octamers, histone H1, 10-base pair arm DNA segments and linker DNA have, however, not yet been elucidated. In the current report, we examine correlations among constituents of nucleosomes from the mechanistic perspectives and present molecular pathways for elucidating supercoiling behaviors of their component DNA sequences. It is our hope that our new analyses could serve as incentives to further clarify correlations between histones and DNA in the dynamic structures of chromatins in the future.

## 1: Introduction

It was demonstrated forty years ago that digestion of eukaryotic chromatins using micrococcal nucleases led to generation of stable complexes of (i) 166 base pairs of DNA, (ii) histone octamers, and (iii) histone H1^1-3^, which are defined as chromatosomes these days^4-6^. Within a structure of chromatosome, 10-base pair DNA sequences at the two ends of chromatosomal DNA are named two 10-base pair arm DNA segments while the rest 146-base pair DNA sequence is called nucleosome core particle DNA (Appendix 1).^7-8^ Structural complexes of chromatosomes along with their covalently connected linker DNA segments are taken as nucleosomes, whose DNA components range commonly from 160 to 240 base pairs in length^9^. These nucleosomal structures are known nowadays as fundamental repeating units of chromatin and chromosome structures of eukaryotic cells.^10-11^

In a series of recent studies of ours, on the other hand, we discovered for the first time that upon binding of histone H1 to polynucleosomes, their naked linker DNA segments turned into negative supercoils^12-15^ (Fig. 1), which determine predominately three-dimensional organizations of chromosomes and chromatins in eukaryotic cells^16^. In spite of the facts that these discoveries were made three years ago^12-15^, detailed interacting patterns among histone octamers, histone H1, 10-base pair arm DNA segments and linker DNA within nucleosomes and chromatins at the molecular scales have not yet been elucidated. In the current report, we examine correlations among constituents of nucleosomes from the mechanistic perspectives and present molecular pathways for elucidating supercoiling behaviors of DNA sequences in nucleosomes and chromatins.

**Fig. 1.**
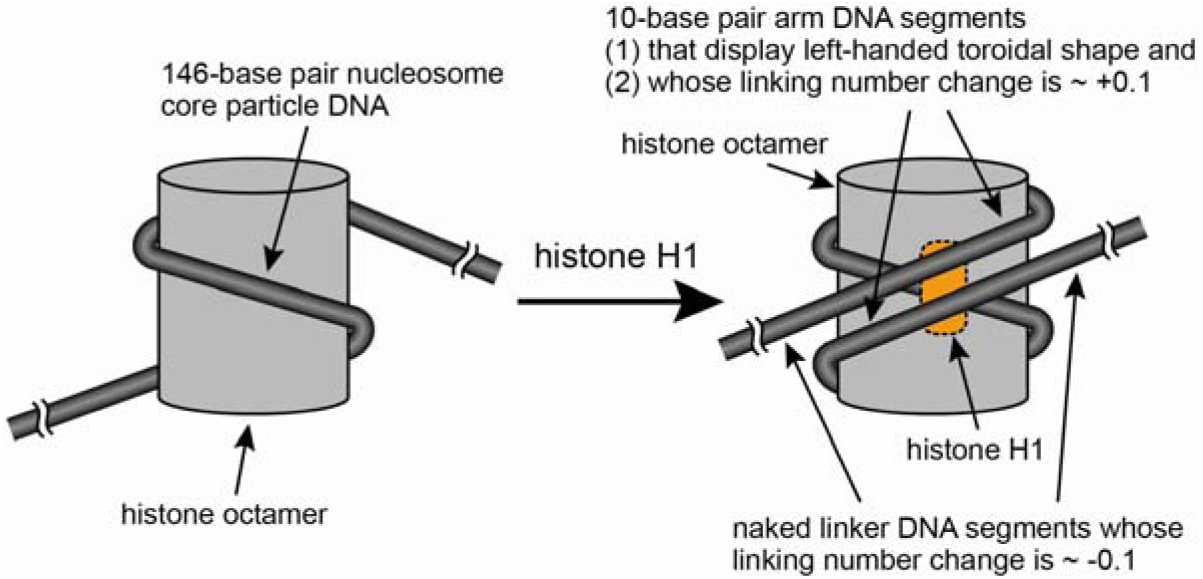
Illustration of our discoveries in the recent years about supercoiling properties of nucleosomes, histone H1, 10-base pair arm DNA segments and naked linker DNA segments^12-16^.

## 2: Physical causes of right-handed alignments of 10-base pair arm DNA segments and their interacting patterns with histone octamers

### 2.1. Complete Contact Model and Incomplete Contact Model of 10-base pair arm DNA segments in their interactions with histone octamers in nucleosomes

In theory, 10-base pair arm DNA segments are able to align along surfaces of histone octamers in two different fashions as illustrated in Complete Contact Model and Incomplete Contact Model in Fig. 2 respectively. In the complete contact model, all of the 10-base pair arm DNA segments are in close physical contacts with surfaces of histone octamers. Because outer surfaces of histone octamers are curved^17-18^, 10-base pair arm DNA segments in the complete contact model must bend themselves in order to achieve complete contact with the surfaces of histone octamers. Extra energy will therefore be required for bending DNA because of high rigidness of double helical structures of DNA^19^ (Fig. 2a).

**Fig. 2.**
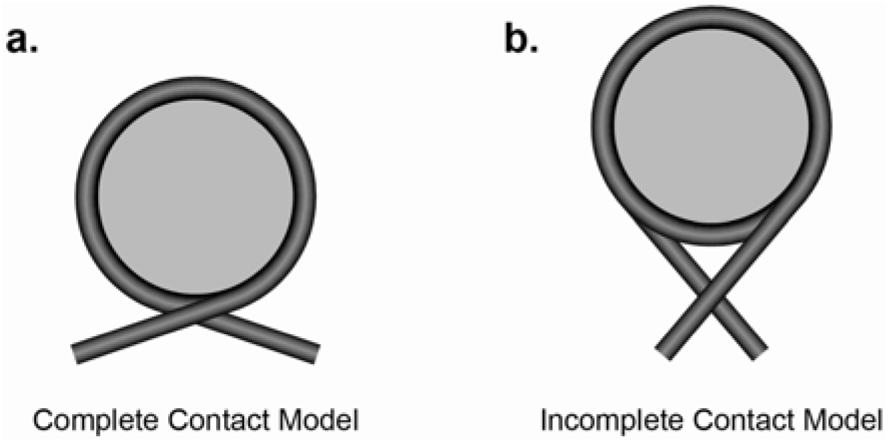
Illustration of Complete Contact Model (a) and Incomplete Contact Model (b) for portraying interacting patterns between 10-base pair arm DNA segments and histone octamers in nucleosomes

In the incomplete contact model (Fig. 2b), on the other hand, not all of the 10-base pair arm DNA segments are in close physical contacts with surfaces of histone octamers. More specifically, some of the 10-base pair arm DNA segments adjacent to ends of 146 nucleosome core particle DNA form complex structures with surfaces of histone octamers while the other base pairs are detached from histone octamers. In the incomplete contact model, 10-base pair arm DNA segments exist in their fairly straight rigid forms. As a result, little extra energy is required to force DNA segments to form curvatures in their duplex backbones. In addition, because backbones of 10-base pair arm DNA segments do not obviously bend back onto surfaces of histone octamer, a cavity space between 10-base pair arm DNA segments and histone octamer exists in the incomplete contact model (Fig. 2b).

### 2.2 Causes of right-handed configuration of 10-base pair arm DNA segments in chromatosomes

Our previous FRET studies showed that when complexes of histone octamers and 166 base pairs duplex DNA were incubated in the presence of histone H1 or ATP, two 10-base pair DNA segments in formed chromatosome displayed were oriented toward each other instead of turning away from each other (Fig. 1). These orientations signify that two 10-base pair DNA segments adopt right-handed toroidal shapes^14^, which make themselves positive supercoils^16^.

In theory, three types of causes that could drive two 10-base pair DNA segments to adopt their right-handed configurations: (1) right-handed alignments of positive charges and polar groups on surfaces of histone octamers (Model 1) and (2) formation of protein-DNA complexes between tails of histones from histone octamers and 10-base pair arm DNA segments (Model 2) and (3) formation of protein-DNA complexes between histone H1 and 10-base pair arm DNA segments (Model 3), which are outlined in the three sections below.

#### (1) Model 1 (histone octamers’ surface-determining model)

Because DNA possesses (1) negatively charged phosphodiester backbones and (2) polar groups, we reason now that positive charges and polar groups could be present and align on the surfaces of histone octamers in a right-handed fashion (Fig. 3a). The (1) electrostatic interactions and (2) hydrogen bonds between (i) 10-base pair arm DNA segments and (ii) surfaces of histone octamers result in formation of relatively stable DNA-protein complexes (Fig. 3b). As a result, the preexistence of right-handed alignment of positive charges and polar groups on surfaces of histone octamers would be able to compel 10-base pair arm DNA segments to adopt their right-handed configurations, which could lead to the observation reported in our earlier studies^14^.

**Fig. 3.**
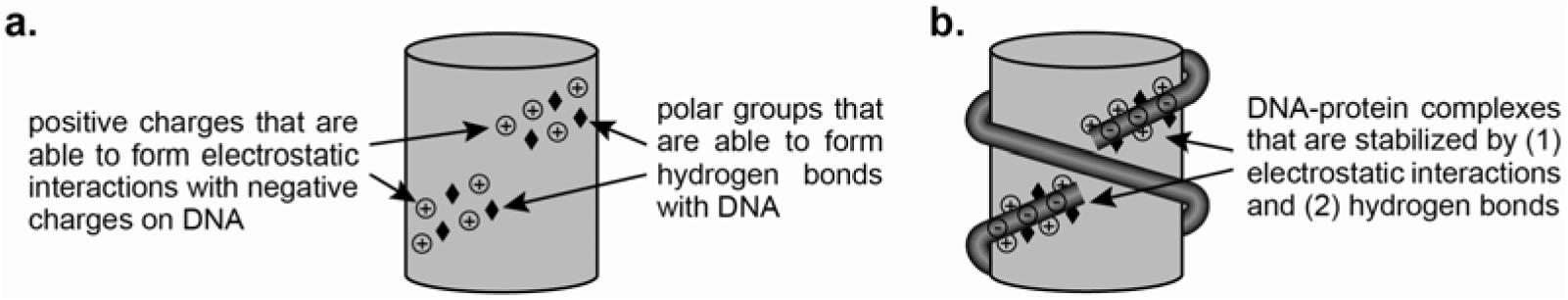
Illustration of (a) right-handed alignment of collective positive charges and polar groups on surfaces of a histone octamer, and (b) formations of DNA-protein complexes between 10-base pair DNA segments and histone octamer through electrostatic interactions and hydrogen bonds, which compel 10-base pair arm DNA segments to adopt right-handed toroidal shapes.

#### (2) Model 2 (histone octamers’ tail model)

It has been known that unstructured tails of histones from histone octamers are present in nucleosome structures.^17-18^ The tails from histone H3, for example, are capable of interacting with DNA segments beyond 146-base pair nucleosome core particle DNA.^20-21^ Consequently, histone octamers will be able to pull 10-base pair arm DNA segments toward themselves through their tails, which could drive arm DNA segments to adopt right-handed toroidal shapes (Fig. 4). In such a model, tails of histone H3 would be able to bind to minor groves of double helical structures of DNA instead of its major groves as various types of peptides do. In addition, it is likely that these minor bindings could lead to overwinding of double helical structures of 10-base pair arm DNA segments, a type of effect that was observable previously from the actions of other known minor groove binders^22-23^.

**Fig. 4.**
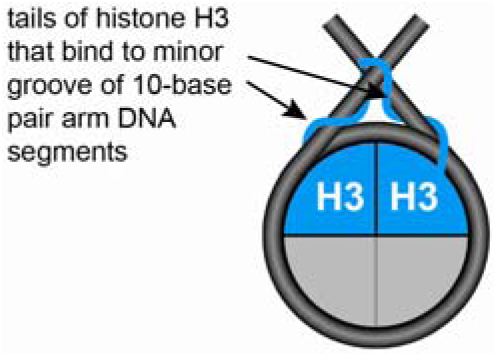
Illustration of formation of DNA-protein complexes between tails of H3 and 10-base pair DNA segments, which forces the adoption of right-handed toroidal shapes by 10-base pair arm DNA segments.

#### (3) Model 3 (histone H1-driven model)

The peptide tails of histone H1 are known to be positively charged.^24-25^ Once this linker histone recognizes and binds to the site adjacent to dyad of a nucleosome (Fi. 5b), its positively charged tails could attach to minor grooves of 10-base pair arm DNA segments (Fig. 5c and Fig 5d), which would be able to neutralize negatively charged phosphodiester DNA backbones for a certain extent. This neutralization could in turn reduce the repelling forces between two separate strands of DNA, thus facilitating approaches of two 10-base pair arm DNA segments to each other. As a result, two 10-base pair arm DNA segments are oriented inward to display right-handed toroidal shapes (Fig. 4).

**Fig. 5.**
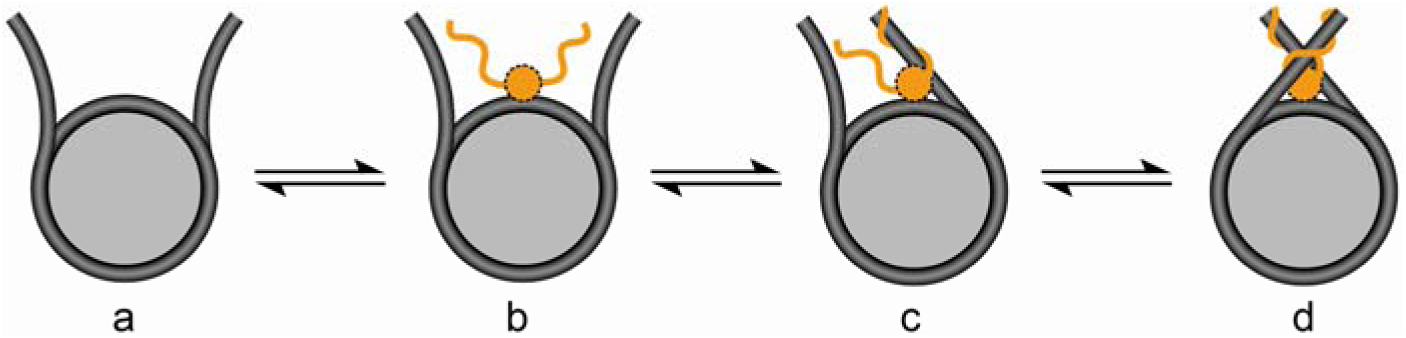
Illustration of formation of right-handed toroidal shapes of 10-base pair arm DNA segments that is driven by actions of (i) histone H1 through its (i) binding to dyad regions and (ii) tails of histone H1 through their bindings to 10-base pair arm DNA segments.

In a brief summary, three models are suggested in the current report for portraying the likely causes of left-handed toroidal shapes of 10-base pair arm DNA segments in chromatosomes. It is anticipated that further studies using Cryo-EM, NMR and x-ray crystallography will be able to provide detailed information about interacting patterns of 10-base pair arm DNA segments with histone proteins in the near future.

## 3. Equilibrium between arm-closed forms and arm-open forms of 10-based pair arm DNA segments in the absence of histone H1

Our previous studies proved that 10-base pair arm DNA segments displayed their arm-open forms and arm-closed forms in nucleosomes respectively in the absence of histone H1.^14^ These observations signify that an equilibrium exists between arm-closed form and arm-open forms in liquid phase and arm-closed form and arm-open forms are two types of stable conformations^14,16^ of 10-base pair arm DNA segments. It was demonstrated in the past that extended micrococcal nuclease’s digestion led to degradation of 10-base pair arm DNA segments from nucleosomes in chromatins.^1-3^ These earlier observations support the notion that in the absence of influence of external agents, the aforementioned equilibrium favors formation of arm-open forms of 10-base pair arm DNA segments (Fig. 6a). We therefore infer that arm-open forms of 10-base pair arm DNA segments are more thermodynamically stable than their arm-closed counterparts under the physiological conditions.

**Fig. 6.**
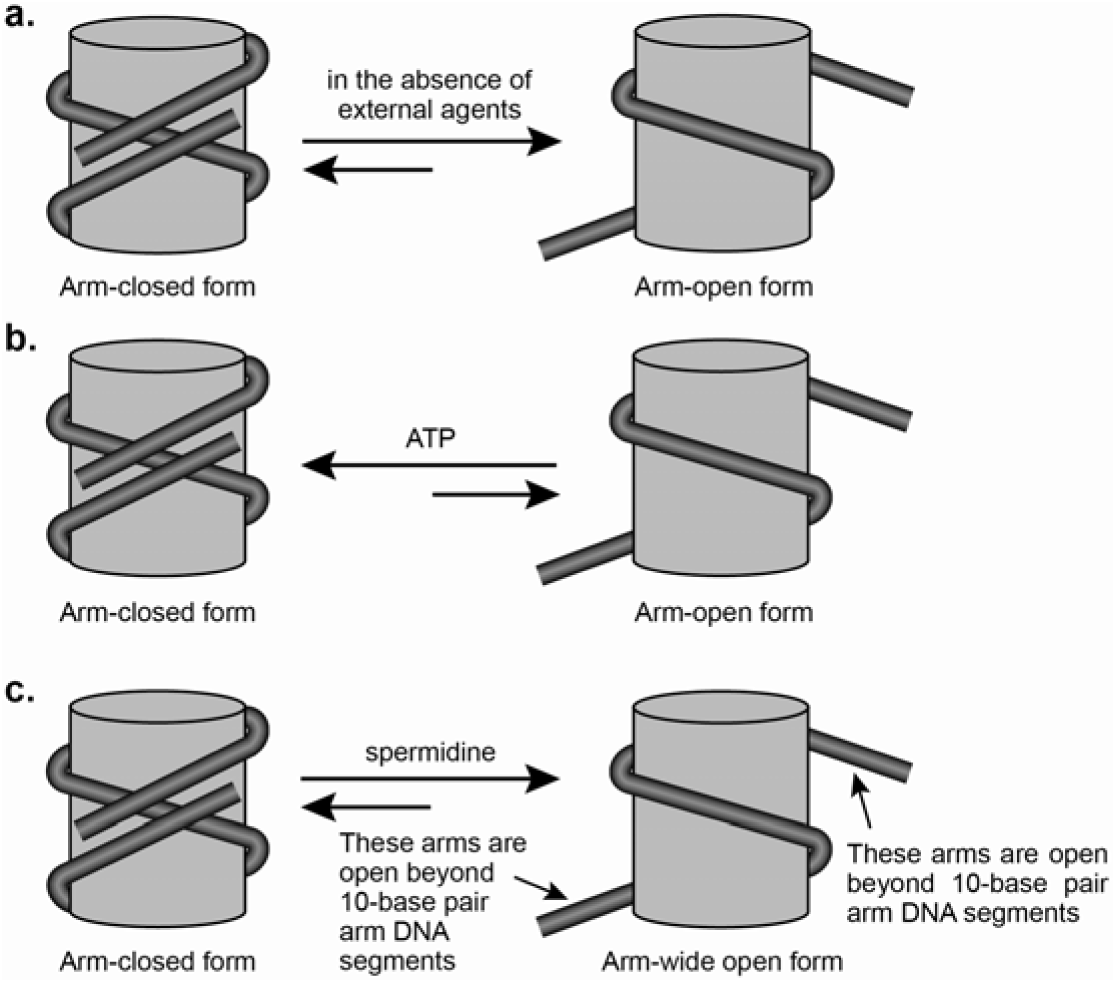
Equilibrium between (a) arm-closed and arm-open forms of 10-base pair arm DNA segments in the presence of neither ATP nor spermidine, (b) in the presence of ATP and (c) in the presence of spermidine.

ATP (adenosine triphosphate), on the other hand, is a type of polyanions and occurs naturally in eukaryotic cells.^26^ Our previous studies showed that in the presence of ATP, nucleosome exhibited predominately their arm-closed forms^14^, which implies that this polycation shifted 10-base pair arm DNA segments’ equilibrium toward their arm-closed forms (Fig. 6b). It is our rationalization that negative charges in ATP are able to reduce the effect of cations that bind to DNA, thus facilitating negatively charged 10-base pair arm DNA segments in their bindings to positively charged surface groups of histone octamers. In addition to ATP^14^, it was showed in our previous studies that spermidine as a polycation forced 10-base pair arm DNA segments to adopt their arm-open forms^14^, which signifies that this polycation shifts 10-base pair arm DNA segments’ equilibrium toward their arm-open forms (Fig. 6c). Since unlike monocations, polycations are able to bind to polyanions through their collective actions of multiple charges, we therefore argue now that tight interactions of spermidine with polyanions of DNA phosphodiester bonds prevent formation of stable complexes between 10-base pair arm DNA segments and histone octamers. As a result, in the presence of spermidine, 10-base pair arm DNA segments are forced to adopt their arm-open forms^14,16^ (Fig. 6c).

To brief sum up, arm-open and arm-closed forms of 10-base pair arm DNA segments are in effect two types of conformations^14,16^ of 10-base pair arm DNA segments, equilibrium of which favors formation of their arm-open forms in the absence of influence of external factors. However, equilibrium constants and rate constants in conversions between arm-open and arm-closed forms can be affected by various external factors such as histone H1, poly-cations (e.g. spermidine), poly-anions (e.g. ATP), temperatures and pH of buffer solutions.

## 4: Roles and pathways of histone H1 in formation of chromatosome structures

### 4.1. Residing positions of histone H1 in a chromatosomal structure

In theory, there are two possible fashions for histone H1 protein to reside in the structural entity of a chromatosome (Model 4 and Model 5 in Fig. 7). In Model 4, histone H1 sits between 10-base pair arm DNA segments and histone octamer (Fig. 7a) while in Model 5, this linker histone binds to 10-base pair arm segments from the DNA side that is opposite to histone octamer (Fig. 7b). It was demonstrated in the past that initial steps of micrococcal nuclease-based digestion of chromatins resulted in leftover of DNA segments of 166 base pairs in length in nucleosomes.^1^ We therefore believe that observations of 166 base pair DNA from micrococcal nuclease digestion evidence that histone H1 is located between histone octamers and 10-base pair arm DNA segments (Model 4 in Fig. 7a). In other words, if histone H1 binds 10-base pair arm segments on the opposite DNA site to histone octamers (Model 5 in Fig. 7b), this linker histone will protect more than 166 base pairs during micrococcal nuclease’s digestion owing to occupancy of its own three-dimensional structures on linker DNA regions. The observed fact was that only 166-base pair DNA sequences resisted micrococcal nuclease’s initial hydrolysis^1^. Consequently, Model 5 might not represent accurate patterns for portraying position of histone H1 in nucleosomes (Fig. 7b). In addition, previous structural analyses on the basis of, NMR^27-29^, electrophoresis^30^ and Cryo EM^8,31^ showed that histone H1 positioned itself near dyad point of nucleosomes inside the arms of 10-base pair arm DNA segments, which is coherent with position of histone H1 in chromatosomes as shown in Model 4 (Fig. 7a).

**Fig. 7.**
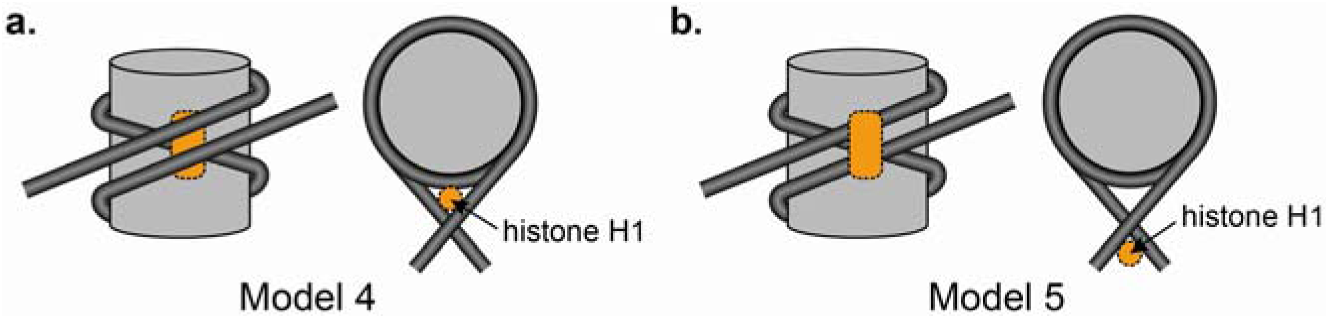
Illustration of two different models (Model 4 (a) and Model 5 (b)) for portraying residing positions of histone H1 within chromatosomes.

It is therefore our conclusion that within the structure of a chromatosome, histone H1 emerges between 10-base pair arm DNA segments and histone octamers near dyad^8,27-31^, which are stabilized mainly through this linker histone’s physical interactions with (i) 10-base pair arm DNA segments and (ii) DNA segments near dyad of nucleosomes (Model 4 in Fig. 7a).

### 4.2. Equilibrium among chromatosomes, histone H1 and histone H1-depleted chromatosomes

It was demonstrated in the past that (1) initial digestion of chromatins using micrococcal nuclease resulted in leftover of 166 base pair DNA segments in chromatosomes and (2) extended digestion led to removal of 10-base pair arm DNA segments from chromatosomes.^1-3^ These observations can be interpreted ad existence of equilibrium among 10-base pair arm DNA segments, histone H1 and histone H1-depleted chromatosomes that caused the initial resistance of DNA in micrococcal nuclease. In theory, histone H1 is incompetent to take the initiative to pull two faraway 10-base pair arm DNA segments together because it is relatively simple protein in structure^24^ and does not possess molecular motor and machinery functions. We therefore suggest that in their equilibrium processes, histone H1 proteins bind to the (i) DNA segments and (ii) histone octamers near dyad of nucleosomes (Fig. 8b) and further wait for incoming 10-base pair arm DNA segments (Fig. 8c). Once 10-base pair arm DNA segments form complexes with histone octamer through (i) electrostatic interactions and (ii) hydrogen bonding, histone H1 seizes the DNA segments through its new (i) electrostatic interactions and (ii) hydrogen bonding, as illustrated in Figs. 8c and 8d.

**Fig. 8.**
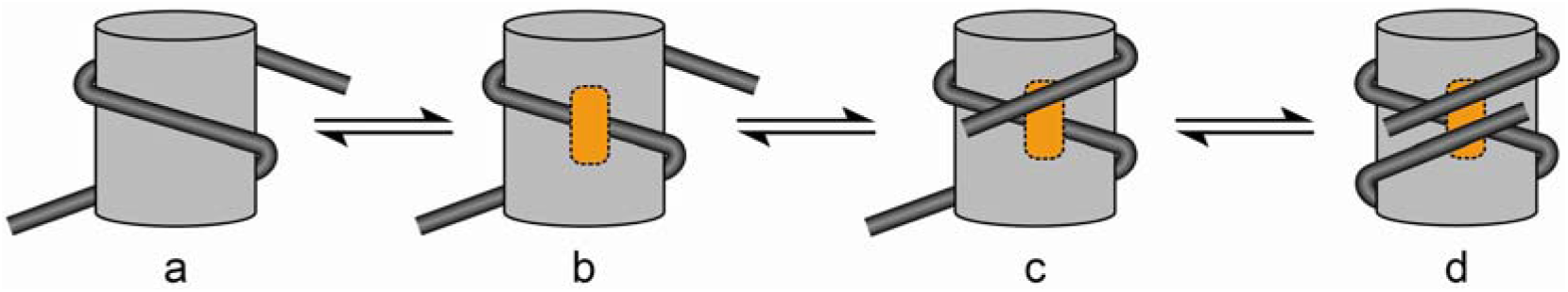
Illustration of equilibrium among histone H1-depleted chromatosomes, 10-base pair arm DNA segments, histone H1 and chromatosomes

Based on our above analysis and reasoning, we infer that four major forms are present in the equilibrium between DNA and histone proteins in their formation of chromatosomes (Fig. 8). Equilibrium constants and rate constants in formation and dissociation of chromatosomes are affected by various external factors such as polycations, poly-anions, temperatures and pH of buffer solutions.

### 4.3. The foremost inborn roles of histone H1 within structures of nucleosome and chromatins

Within the structures of 30-nm chromatin fibers, histone H1 resides within the arms of 10-base pair arm DNA segments^8,27-31^ (Fig. 5a). At first glance, functions of histone H1 in such structures might be (i) to hold 10-base pair arm DNA segments together and (ii) to maintain the structural integrity of chromatosomes. However, when viewed from the supercoiling standpoint, we argue that the foremost inborn role of histone H1 is to serve as a blockage to insulate (i) positively supercoiled 10-base pair arm DNA segments and (ii) negatively supercoiled respectively. More specifically, once histone H1 resides in the middle of (i) 10-base pair arm DNA segments and (ii) naked linker DNA segments, supercoils of opposite signs in the two types of DNA segments cannot be neutralized (Fig. 9a). Such topologically insulating roles of histone H1 can be explicitly illustrated using simplified histone H1-bound circular double helical structures of DNA as shown in Fig. 9b. In such non-histone octamer-containing structures, owing to the binding of histone H1 to two opposite strands of DNA, positive supercoils and negative supercoils are separated and coexist.

**Fig. 9.**
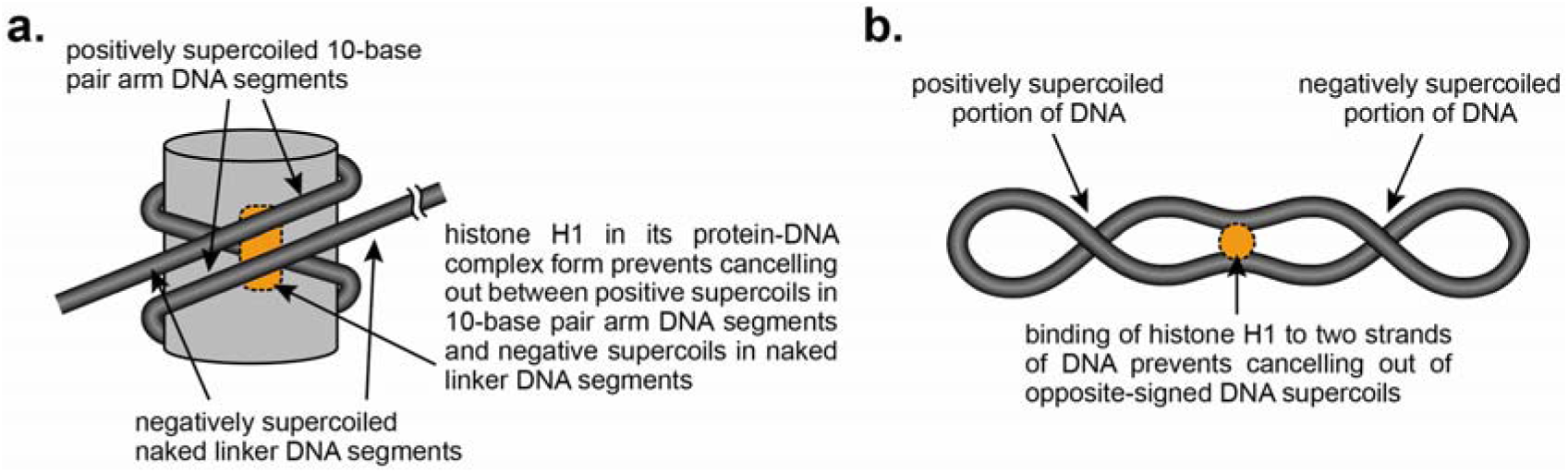
Illustration of (a) histone H1’s blockages of neutralization between positively supercoiled 10-base pair arm DNA segments and negatively supercoiled naked linker DNA segments in chromatins and (b) histone H1’s blockages of neutralization between two opposite signed DNA supercoils, in which histone octamers are not present.

In a brief summary, once histone H1 binds to the stretches near ends of 10-base pair arm DNA segments, it serve mainly as a topological blockage to insulate between (i) positive supercoils in 10-base pair arm DNA segments and (ii) negative supercoils in naked linker DNA regions.

## 5 Supercoils within double helical structures of 10-base pair arm DNA segments

Nucleosome core particle DNA possesses 146 base pairs in length, which wraps around histone octamer for ~1.75 turns.^17-18^ One of the most remarkable physical characteristics of this 146-base pair DNA segments is its possession of both negative supercoils and positive supercoils simultaneously in its structure.^32-33^ The negative supercoils are caused by its adoption of left-handed toroidal overall shapes, which gives rise to linking number change of −1.75.^19^ The positive supercoils, however, are produced within its internal structures as a result of overwinding of its double helix, which leads to linking number change of +0.75.^34-35^ Consequently, the net change of linking number in the 146 base pairs of nucleosome core particle DNA is −1 ((−1.75) + (+0.75) = −1), as observed in experimental examinations of linking number changes in plasmid DNA^36^.

The aforementioned simultaneous adoption of both negative supercoils and positive supercoils in 146 base pair nucleosome core particle DNA^32-36^ raises a question regarding whether 10-base pair arm DNA segments could undertake two types of different forms of supercoils in their double helical structures concurrently as well. Our earlier studies proved that 10-base pair arm DNA segments take on right-handed toroidal shapes on a whole, which are positively supercoiled.^14^ However, the supercoils within double helical structures of 10-base pair arm segments have not yet known experimentally.^14,16^ In theory, there are three possible types of double helical structures that 10-base pair arm DNA segments could adopt: (1) underwound form (negative supercoils) (Fig. 10a), (2) relaxed form (non-supercoils) (Fig. 10b) and (3) overwound form (positive supercoils) (Fig. 10c).

**Fig. 10.**
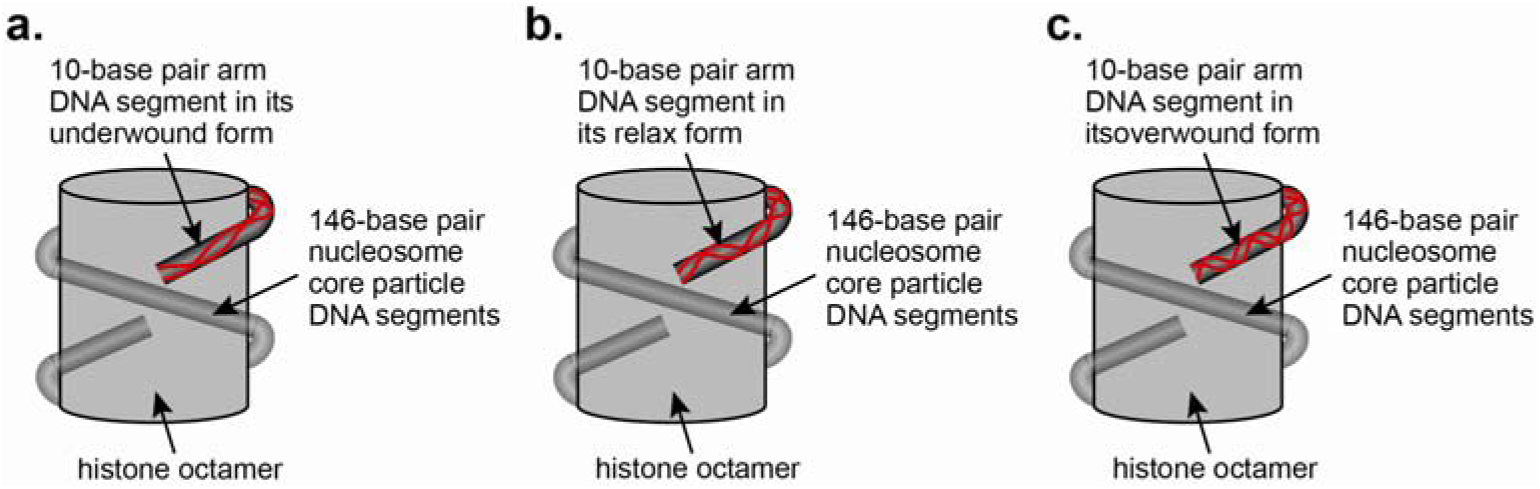
Illustration of double helical structures of 10-base pair arm DNA segments that are in their under-wound (a), relaxed (b), and overwound forms respectively.

If 10-base pair arm DNA segments adopted underwound structures, its resultant negative supercoils would be cancelled out with positive supercoils affiliated with their right-handed alignments along surfaces of histone octamers^14^ to a large extent. Our previous studies showed that magnitudes of negative supercoils in naked linker DNA regions were in a reasonable range^13^, which is the indirect evidence that 10-base pair arm DNA segments unlikely adopt their negatively supercoiled underwound forms. However, there has been lack of experimental evidence and logic foundations for us to make a judgment on which of the additional two types of double helical structures, underwound or relaxed, are actually undertaken by 10-base pair DNA segments in their arm-closed forms. We anticipate that new Cryo-EM-, NMR-based studies will be able to resolve these issues in the near future. On the basis of our above deduction and analyses, we infer that when they are in their arm-closed forms in nucleosomes, double helical structures of 10-base pair arm DNA segments can be either overwound or relaxed, and cannot be underwound.

## 6. Correlations between magnitudes of superhelical densities in naked linker DNA segments and structural forms of polynucleosomes

### 6.1 Choices of structural forms by polynucleosomes in the presence of histone H1

Our earlier studies showed that as a result of binding of one histone H1 to one nucleosome, linking number in its corresponding naked linker DNA regions is altered by ~ −0.09, which means that collective binding of 11 histone H1 proteins to 11 nucleosomes in polynucleosomes leads to change of linking number by ~ −1.^13^ Within a structure of single unit of nucleosome that possess open DNA terminuses, negative supercoiling in their naked linker DNA segments is not preservable. This happens because generated negative supercoils by binding of histone H1 are releasable through open ends of their naked linker DNA segments. In the event of circular polynucleosomes, however, negative supercoils in their naked linker DNA regions cannot be releasable upon binding of histone H1 because there is absent of open end in their circular structures.^37^ Besides circular polynucleosomes, when linear polynucleosomes possess comparably high molecular weights (*e.g.* chromatins), terminuses of their DNA macromolecules are not rotatable freely in practice so that negative supercoils in their naked linker DNA segments can be preserved as well.^16^ These preserved negative DNA supercoils can be described quantitatively using superhelical density (σ)^38^, which is expressed in form of the following equation:

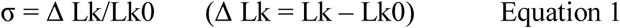

where Lk0 is the linking number of DNA in its relaxed form and Lk is the linking number of DNA in its supercoiling states^38^.

According to our newly formulated supercoiling theory of chromosomes^16^, lengths of linker DNA of polynucleosomes in eukaryotic cells could vary significantly and play critical roles in determining spatial organizations of chromosomes. If lengths of naked linker DNA segments in polynucleosomes are (i) equal to or less than 30 base pairs, (ii) between 30 and 50 base pairs, and (iii) equal or longer than 70 base pairs, they are named (i) densely packed polynucleosomes, (ii) loosely packed polynucleosomes and (iii) slack poly-nucleosomes respectively (Appendix 1).^16^ Upon binding of histone H1, (i) densely packed polynucleosomes and (ii) loosely packed polynucleosomes are capable of turning into (ii) densely packed 30-nm chromatin fiber and (ii) loosely packed 30-nm chromatin fibers respectively (Appendix 1). The causes of transitions from (i) densely packed and (ii) loosely packed polynucleosomes to (i) densely packed and (ii) loosely packed 30-nm chromatin fibers are changes of magnitudes of superhelical densities in their naked linker DNA segments (Fig. 11). More specifically, when naked linker DNA segments in a nucleosome in polynucleosomes possess 30-base pairs in length, Lk_0_ is 2.88 (30 base pairs/10.4 base pairs ≈ 2.88). Upon binding of one histone H1 protein to one nucleosome, linking number of ~ −0.09 will be introduced in to its linker DNA segments, which gives rise to Δ Lk of ~ −0.09. On the basis of assessments using Equation 1, superhelical density (σ) in naked linker DNA segments of 30 base pairs in histone H1-bound polynucleosomes will be −0.03 (σ = −0.03 (σ = −0.09/2.88= −0.03)^38^. Similarly, it can be derived that magnitudes of superhelical density will be −0.02 and −0.01 in naked linker DNA segments of (i) 50-base pairs (loosely packed polynucleosomes) and (ii) 70-base pairs (sack polynucleosomes) in length (Table 1).

**Fig. 11.**
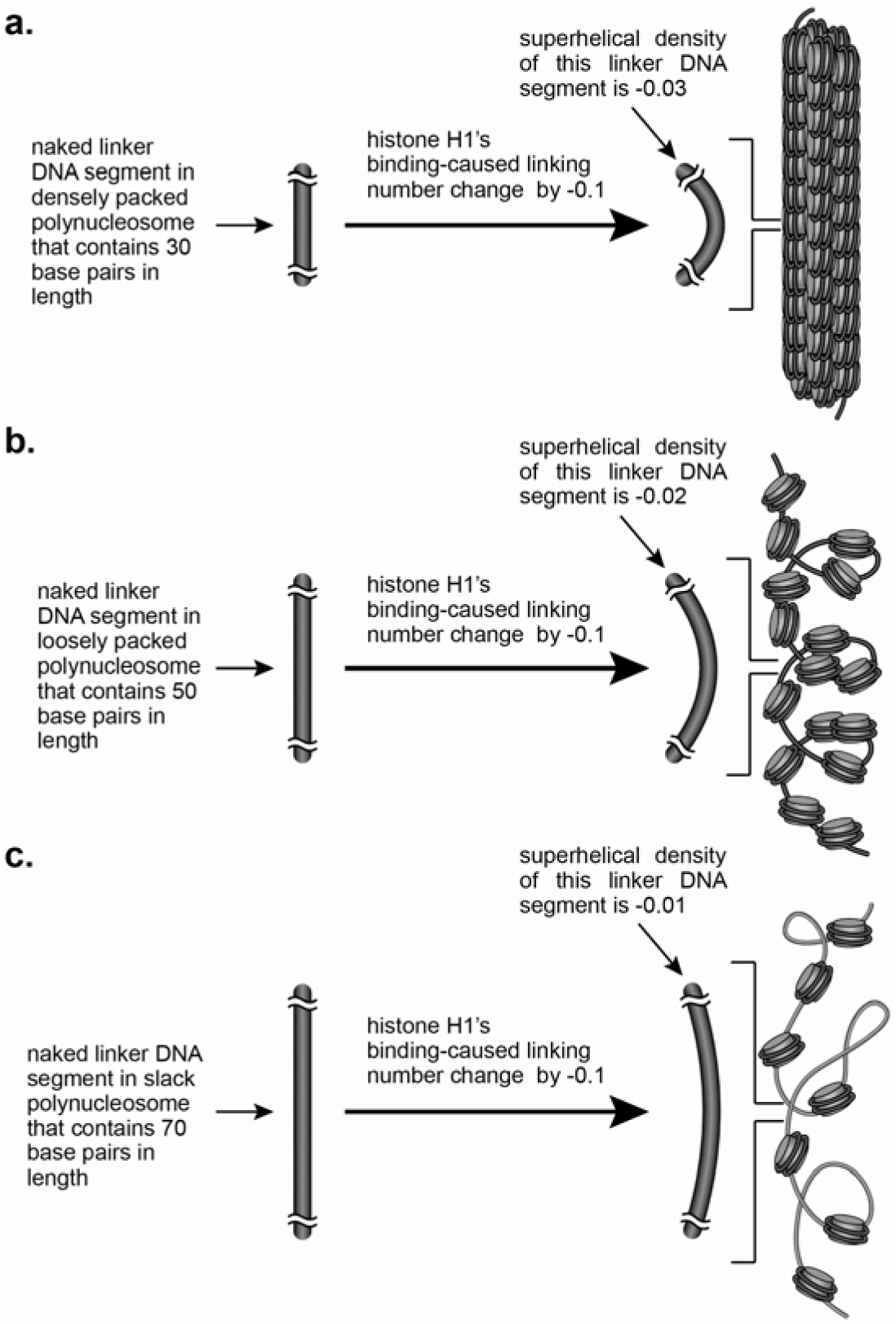
Illustration correlation between superhelical densities in their naked linker DNA segments and structural forms of polynucleosomes. (a) 30 base pairs in naked linker DNA segments of polynucleosomes, (b) 50 base pairs in naked linker DNA segments of polynucleosomes, and (c) 70 base pairs in naked linker DNA segments of polynucleosomes.

From the supercoiling standpoint, on other hand, increase of superhelical density always causes rise of backbone curvatures of double helical structures of DNA^39-40^. We therefore reason that these high superhelical density-driven high degrees of backbone curvatures cause densely packed polynucleosomes to form densely packed 30-nm chromatin fibers (Table 2). As opposed to densely packed polynucleosomes, sack polynucleosomes, however, are not able to form 30-nm chromatin fiber-like structures because superhelical density in their naked linker DNA segments remain significantly low upon binding of histone H1.

**Table 2.**
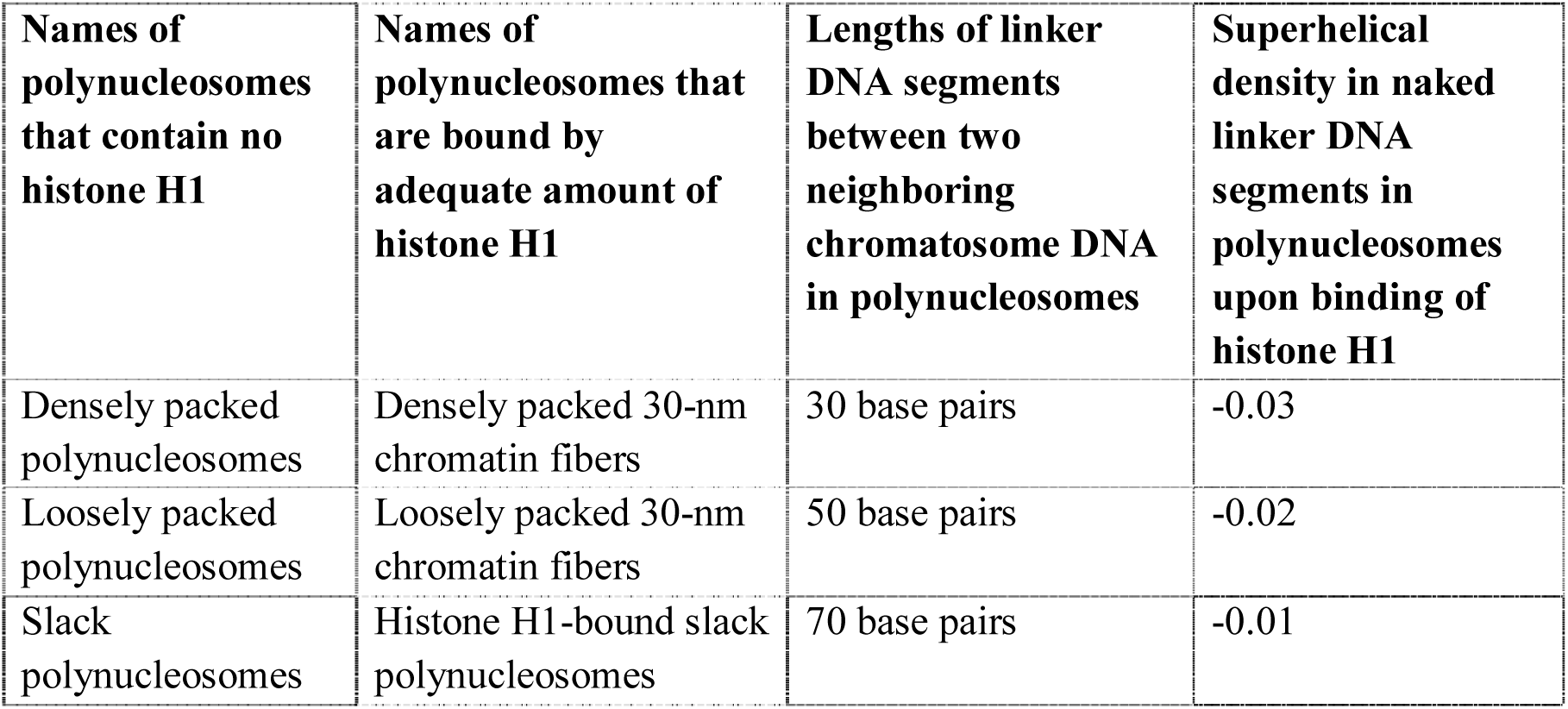
Correlation of lengths of naked linker DNA segments in polynucleosomes with magnitudes of their superhelical densities upon binding of histone H1 proteins.

Based on the above analysis and reasoning, we conclude that magnitudes of superhelical densities in their naked linker DNA segments govern what types of structural forms (densely packed 30-nm chromatin fibers, loosely packed 30-nm chromatin fibers or histone H1-bound slack polynucleosomes) histone H1-bound polynucleosomes will adopt.

### 6.2 Choices of structural forms by polynucleosomes in the (i) presence of ATP and (ii) absence of histone H1

Our previous studies showed that in the (i) presence of ATP and (ii) absence of histone H1, 10-base pair arm DNA segments in single nucleosomes was capable of adopting their arm-closed forms^14^. It is therefore reasonable to suggest that ATP-affiliated arm-closed forms will lead accumulations of negative supercoils in naked linker DNA regions in polynucleosomes to a certain degree as histone H1-forced arm-closed forms do^13,15^. On the basis of our analyses shown in Section 6.1, we conclude that densely packed 30-nm chromatin fibers could form from polynucleosomes in the absence of histone H1 if (1) sufficient high concentrations of ATP are present and (2) lengths of their component naked linker DNA segments are adequately short.

### 6.3 Choices of structural forms by polynucleosomes in the presence of neither histone H1 nor ATP

Existence of equilibrium between arm-open and arm-closed forms in nucleosomes signifies that at any given moment, certain portions of arm-closed forms of 10-base pair arm DNA segments exist in polynucleosomes (Fig. 6a). We therefore rationalize that negative supercoils can be accumulated and preserved in naked linker DNA regions of polynucleosomes to a low extent in the absence of histone H1 and ATP. As long as lengths of naked linker DNA in polynucleosomes are adequately short, high superhelical densities can be generated, which will lead to formation of densely packed 30-nm chromatin fibers (Fig. 12). However, 30-nm fibers cannot form from polynucleosomes when lengths of their component linker DNA segments are comparably long. Even though 30-nm fibers cannot form from the polynucleosomes whose naked linker DNA segments possess relatively long length, these polynucleosomes will still hold left-handed toroidal shapes owing to the presence of negative supercoils in their naked linker DNA segments. These negative supercoils are affiliated with the arm-closed forms (Fig. 6b) that are in equilibrium with their counterparts of arm-open forms (Fig. 6a),

## 7. Differences between negative supercoils in naked linker DNA regions of polynucleosomes and those produced by enzymatic actions

The processes of generations of DNA supercoils and their relaxations in living cells can be classified into the following three categories:

### (1) Category 1

*Accomplishment of generation and relaxation of DNA supercoils by enzymatic activity*. Examples of enzymes in this category are (i) gyrase and prokaryotic topoisomerase I in prokaryotic cells as well as (ii) reverse gyrase and hyperthermophilic topoisomerase I in hyperthermophilic bacteria;

### (2) Category 2

*Accomplishment of generation and relaxations of DNA supercoils by combined physical actions of proteins and chemical actions of enzymes*. Examples in this category are actions of pairs of (i) helicase and topo I (or topo II) as well as (ii) histone octamers in the formation of nucleosome core particles and topo I (or topo II); and

### (3) Category 3

*Accomplishment of generations and relaxation of DNA supercoils merely by physical association of proteins with and their dissociation from DNA without involvement of chemical reactions.* Examples in this category are (i) association of 10-base pair arm DNA segments with histone octamers and their dissociations from histone octamers^14^ as well as (ii) binding of Condensin I protein complexes to metaphase chromosomal DNA and their dissociation from chromosomes^41-42^.

In Category 1 and Category 2, chemical actions of enzymes preserve altered supercoiling states (linking number changes) in DNA macromolecules in the later stages of their activities^38,43^. These preservations mean that after enzymes are dissociated from DNA, altered linking numbers remain and can hence be detected subsequently using gel electrophoresis^44^ or by other physical means such as AFM^45^. In Category 3, however, interacting patterns between proteins and DNA segments fundamentally differ from those in Category 1 and Category 2. For example, linking number changes caused by interactions between 10-base pair arm DNA segments and histone octamers are merely physical processes, which are never chemically preserved. That is, when 10-base pair arm DNA segments are detached from histone octamers and histone H1, altered linking numbers at its initial step is neutralized with naked linker DNA segments. As a result, physically reversible DNA supercoils in Category 3 have often gone unnoticeably during experimental examinations in the past^8,26-28,30,45-46^.

In addition, in the event of gyrase (Category 1) and topo II (Category 2), actions of these enzymes always lead to changes of linking number by |1| or more^37,42^. When 10-base pair arm DNA segments bind to a histone octamer, however, it will generate linking number change by |0.09|^13^. This significantly low magnitude of linking number changes have often been unidentifiable under the examinations of x-ray crystallography^46-47^, NMR^27-29^ and Cryo-EM^8,31^.

## 8. Conclusion

It was demonstrated in our previous studies that 10-base pair arm DNA segments displayed positively supercoiled right-handed toroidal shapes.^14^ Our current analyses show that right-handed configurations of these 10-base pair arm DNA segments are in effect determined by structures of histone proteins, as it happens to 146-base pair nucleosome core particle DNA^19^. In addition, different from those produced by enzymatic actions^38,43^, generation of supercoils in 10-base pair arm DNA segments and naked linker DNA regions are types of physical processes and easily reversible. This easy reversibility in supercoiling alterations is constantly associated with arm-open form of nucleosomes, arm-closed forms of nucleosomes, histone H1 and chromatosomes, which are in dynamic equilibrium in a liquid phase. In spite of the fact that various types of structural data about nucleosome and chromatins have been acquired to date^8,27-29,31,46-47^, however, certain critical questions remains unsolved such as supercoils in the internal double helices of 10-base pair arm DNA segments as well as holding patterns of 10-base pair arm DNA segments by histone proteins. It is anticipated that resolution of these uncertain issues will rely on further x-ray crystallography-, NMR- and Cryo EM-based examinations on nucleosomes and chromatins in the near future.

## APPENDIX 1 Technical terms and axioms used for describing supercoiling properties of DNA

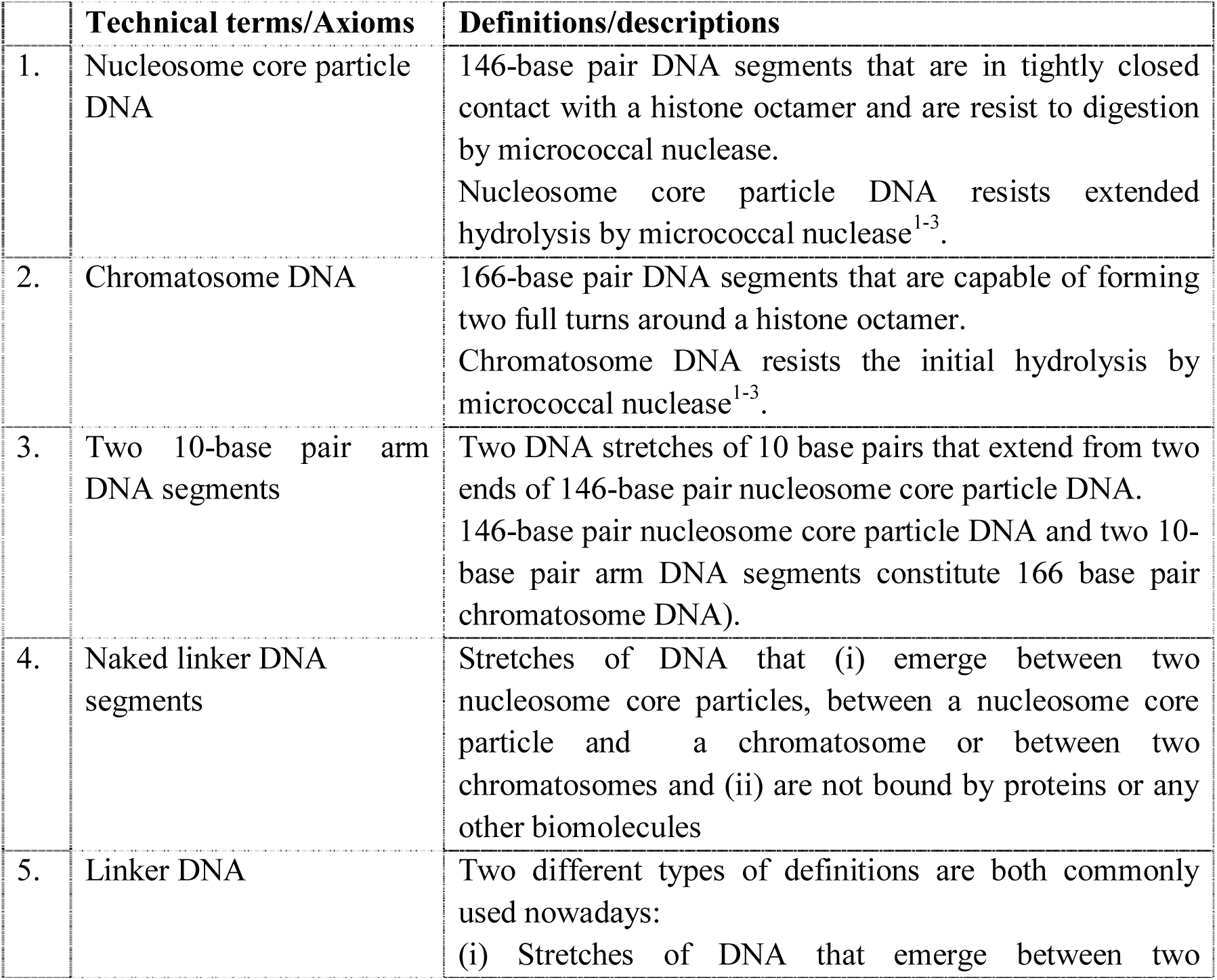
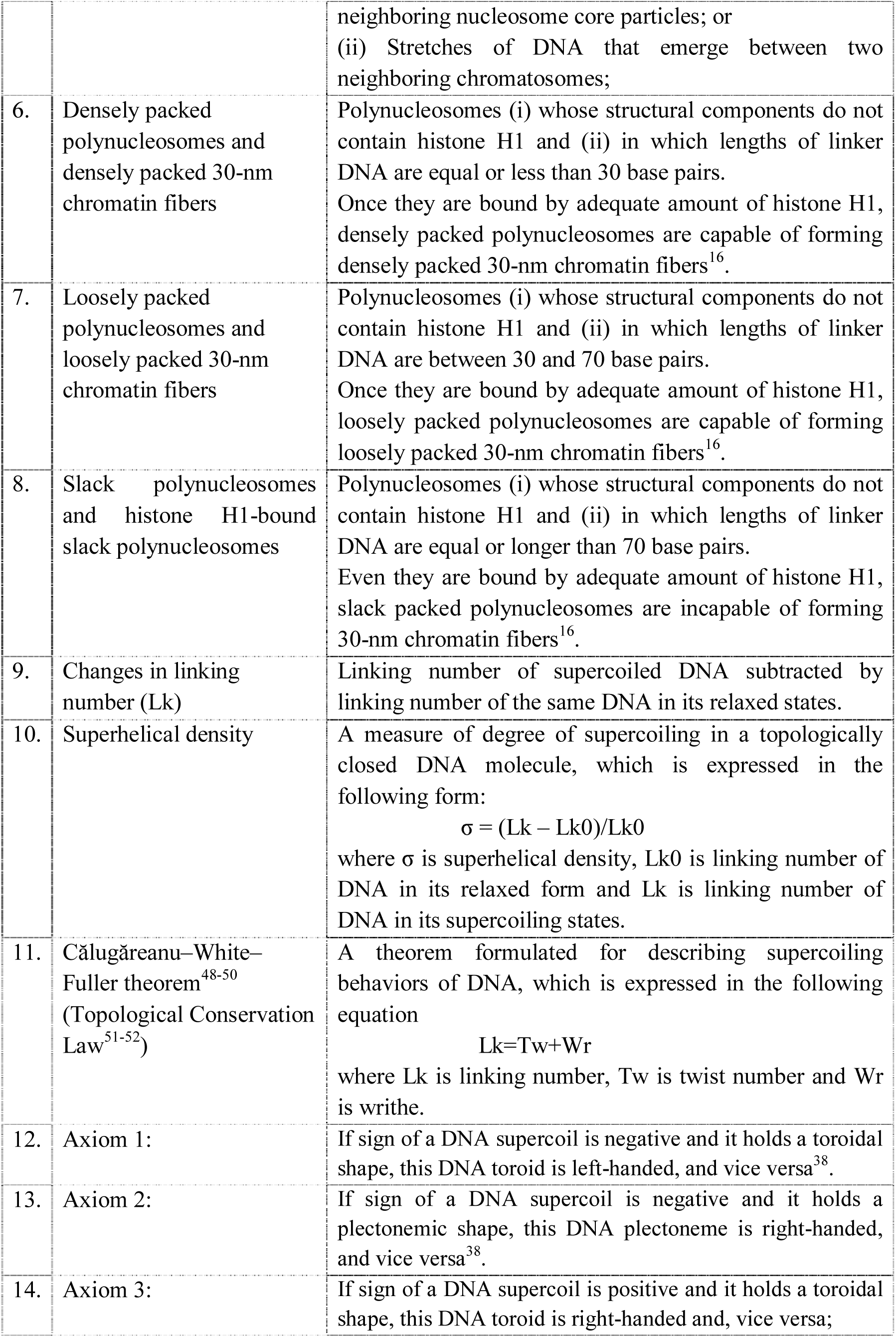
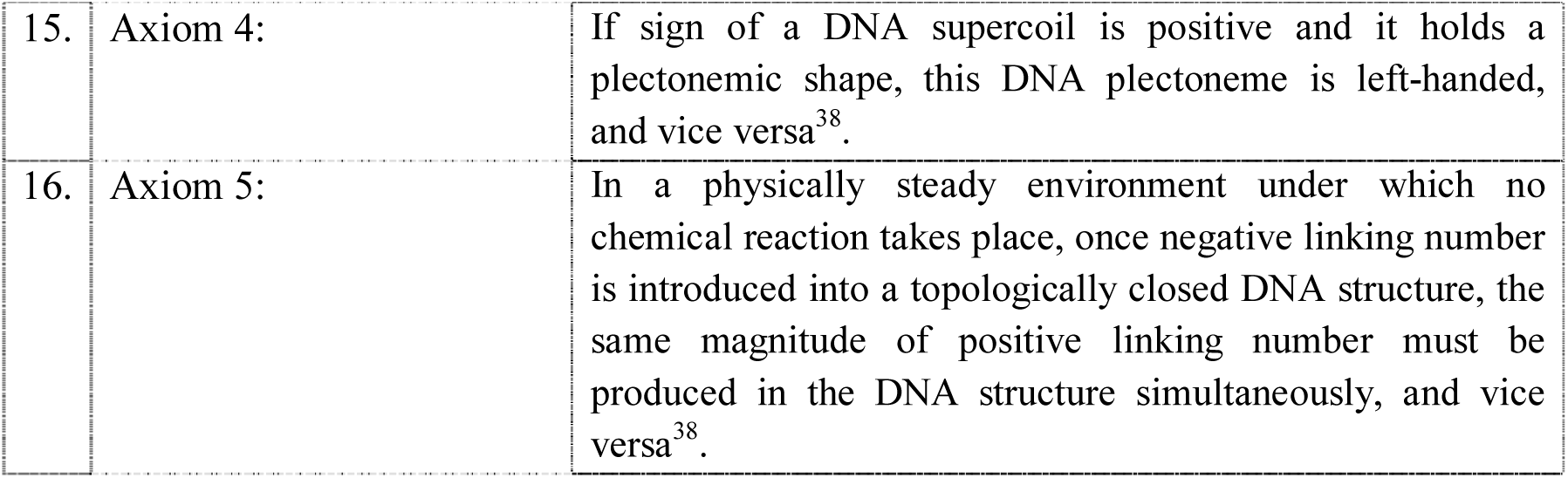
Related to Methods. Summary statistics for RNA-seq samples alignments.

## Acknowledgement

This work was supported by Ministry of Education in Singapore, Nanyang Technological University and Agency for Science, Technology and Research in Singapore through research Grants to Tianhu Li (MOE2014-T2-2-042, MOE RG14/15, MOE RG13/16, MOE RG117/17 and SERC A1883c0007).

